# Simulation-based dosimetry of transcranial and intranasal photobiomodulation of the human brain: the roles of wavelength, power density and skin colour

**DOI:** 10.1101/2024.04.05.588330

**Authors:** Hannah Van Lankveld, Anh Q. Mai, Lew Lim, Nazanin Hosseinkhah, Paolo Cassano, J. Jean Chen

## Abstract

Photobiomodulation (PBM) is a novel technique that is actively studied for neuromodulation. However, despite the many in vivo studies, the stimulation protocols for PBM vary amongst studies, and the current understanding of neuromodulation via PBM is limited in terms of the extent of light penetration into the brain and its dosage dependence. Moreover, as near-infrared light can be absorbed by melanin in the skin, skin tone is a highly relevant but under-studied variable of interest. In this study, to address these gaps, we use Monte Carlo simulations (with MCX) of a single laser source for transcranial (tPBM) and intranasal (iPBM, nostril position) irradiated on a healthy human brain model. We investigate wavelengths of 670, 810 and 1064 nm in combination with light (“Caucasian”), medium (“Asian”) and dark (“African”) skin tones. Our simulations show that a maximum of 15% of the incidental energy for tPBM and 1% for iPBM reaches the cortex from the light source at the skin level. The rostral dorsal prefrontal cortex in tPBM and the ventromedial prefrontal cortex for iPBM accumulates the highest highest light energy, respectively for both wavelengths. Specifically, the 810 nm wavelength for tPBM and 1064 nm wavelength for iPBM produced the highest energy accumulation. Optical power density was found to be linearly correlated with energy. Moreover, we show that “Caucasian” skin allows the accumulation of higher light energy than other two skin colours. This study is the first to account for skin colour as a PBM dosing consideration, and provides evidence for hypothesis generation in in vivo studies of PBM.

## 1. Introduction

Transcranial photobiomodulation (tPBM) is defined as the application of low levels of red or near-infrared light (NIR) to stimulate neural tissue [1]. The act of shining light on the head to modulate brain function has been utilized in the field of neuroscience since 1967 [2]. Originally referred to as low-level laser therapy (LLLT), the incoming light was used for wound healing and pain reduction. NIR light can penetrate human skin and tissue to various depths of several millimetres [1]. This penetration allows the light to stimulate tissue at an intracellular level. It has been proven that absorbing light can give rise to biological reactions of the living system [3]. Light can directly excite photoreceptors in cells, including neurons and endothelial cells. In the NIR or red spectrum, the main chromophore is cytochrome c oxidase (CCO), which is the terminal respiratory-chain enzyme (complex IV) located in the cellular mitochondria. NIR or red light irradiation of the mitochondria can induce the release of nitric oxide (NO), increasing CCO metabolism and leading to the elevation in adenosine triphosphate (ATP) production [4], [5], [6], [7]. While ATP production can in theory enhance mitochondrial function, NO release can lead to vasodilation and increase cerebral blood flow (CBF).

Over the next 50 years LLLT research gained momentum, and more recently was recharacterized as photobiomodulation. Light-emitting diodes (LEDs) were progressively added into the fold, especially in commercial use. LEDs are more easily accessible and cheaper than the original laser source. However, although more convenient, LEDs produce a broader spectrum of light, generating a less focused beam and scatters more when contacting the multi-layer tissue of the human head.

The type of light source (LED vs. laser) is just one variable that impacts the overall effectiveness of PBM. To effectively penetrate the human head’s multi-layered tissue with a beam of light, several parameters can be optimized: (1) wavelength (*λ*), (2) optical power density (OPd), (3) light pulsation frequency (*f*), (4) irradiation duration (*T*), (5) light source positioning. A change in each parameter can dictate a different outcome of PBM application. Previous research, based on cultured cortical neurons, suggests that the peak PBM response happens when the energy induced by the light source (i.e. the fluence) reaches 0.3-3 J/cm^2^ in the specified brain region [8]. Each parameter can be adjusted to impact the overall energy accumulation in tissue. The range of parameter values used in past studies is summarized in **Table 1**.

**Table 1.** Past research parameter values [9], [10], [11].

| PARAMETERS OF INTEREST | UNIT | COMMON RESEARCH VALUES |
| --- | --- | --- |
| Wavelength | $\lambda$ | 532 - 1070 nm |
| Optical Power Density | OPd | 0.1 - 1000mW/cm<sup>2</sup> |
| Pulsation Frequency | $f$ | 1 - 10,000Hz |
| Duration | $T$ | 1 - 20 minutes |

As shown in **Table 1**, the choice of these values is highly variable. A recent 2019 review paper demonstrated the pro-cognitive effects of tPBM in healthy young adults and reported a large difference in selected stimulation parameters; OPd range: 44.4-285 mW/cm2 and λ: 633 nm, 850 nm and 1064 nm [10]. While the majority of PBM studies are transcranial (tPBM), more recently, additional areas of light source positioning have been studied, including intranasal stimulation (iPBM) [9]. In the context of this work, we will use tPBM to refer to scenarios in which the light needs to penetrate the cranial bone, and iPBM to refer to scenarios in which the light travels through the porous cribriform plate.

Despite this vast range of parameters, cognitive improvements have been reported in all these studies. Specifically, previous work has shown that a 1064nm laser can increase oxidized cytochrome-c-oxidase [12], elevate oxygenated hemoglobin concentrations [13][14] and increase prefrontal vascular oxygenation [7]. For the treatment of Alzheimer’s disease (AD), 810nm LEDs had a positive effect on improving desired brain activity [15] as well as increased cerebral perfusion and connectivity [16]. In mouse models, 635nm showed that tPBM could improve brain microenvironmental conditions and alleviate cognitive deficits [17]. 660nm alleviated cognitive dysfunction and reduced microgliosis [18], and 1070 nm triggered microglia responses and promoted angiogenesis [19]. For the treatment of depression, PBM has had promising results with a vast range of parameters. An 823 nm LED, with 36 mW/cm^2^ power density demonstrated antidepressant properties in patients with Major Depressive Disorder (MDD) [20]. In 2020, 945 nm LED improved brain activity and stated that it may clinically decrease anxiety and depression [21]. However, contradicting these findings was a 2022 paper, using an 830 nm LED, that showed no significant differences between tPBM and sham experiments for the treatment of MDD [22]. This study concluded that the optimal dose of tPBM has yet to be defined. Other varied reports are available for a variety of diseases treated with PBM.

Building on promising behavioural outcomes in previous studies, there are approximately 300 registered PBM clinical trials on the effects of PBM on various brain diseases, including Alzheimer’s disease (AD), Parkinson’s disease (PD), depression, back pain, traumatic brain injury (TBI), plantar fasciitis, chronic obstructive pulmonary disease and many more [23]. However, PBM remains a relatively new stimulation method that has many unknowns regarding how much the light interacts with brain tissue [24]. Moreover, despite the large number of on-going studies, there are no universal settings for the tPBM stimulation parameters.

Many researchers rely on historical literature for determining variable values. Selecting suboptimal stimulation parameters can result in diminished PBM responses, potentially hindering progress in PBM research. This highlights the urgent need for research aimed at identifying an optimal set of parameters that maximize energy accumulation in the brain. Previous research has employed a similar model approach to estimate the light penetration into the brain [25], [26], and found that the 810nm wavelength consistently results in the highest energy deposition for tPBM when compared to other frequently employed wavelengths [25]. Moreover, when comparing light source placement in the left and right hemispheres, the values were similar, with the left hemisphere receiving only a marginally higher amount of energy, approximately 1.6% [26]. These previous publications have utilized many LED arrays and optical coefficients without considering the influence of *skin colour*.

This study investigates the effect of melanin on light penetration. Melanin is a natural skin pigment that absorbs light energy upon interaction, with the extent of absorption corresponding to the amount of melanin present. Simultaneously, the skin’s surface can scatter light that remains unabsorbed. This scattering process alters the direction of light as it interacts with specific tissues or mediums. Both scattering and absorption are influenced by melanin content, thereby influencing skin colour. The colour of human skin is influenced by the presence of melanin pigment. The amount and type of melanin in your skin determine its colour. The variation in skin colour among individuals is primarily due to genetic factors. Those with higher melanin levels tend to have darker skin, while those with lower melanin levels have lighter skin. Thus, our study also focuses on understanding the impact of skin colour to the penetration of light and deposition of energy in the human brain. To better understand how each area of the brain is impacted by different light source positioning and stimulation parameters, we simulate the light propagation into the brain at various settings. Our simulations will concentrate on a single-source laser strategically placed between the left and right hemispheres, spanning both common tPBM and iPBM positioning.

## 2 Methods

### 2.1 The Monte Carlo method

The Monte Carlo technique can be utilized to simulate the near-infrared light propagation through the multi-layer tissues, and has been widely used to simulate the effect of PBM in both human and rat brains [25], [26], [27], [28]. When a photon strikes an object, its trajectory is determined by three main properties: (1) absorption, when the neural tissue absorbs the incoming photon, (2) scattering, when the incoming photon is ricocheted and diverted from its original path, and (3) transmission, when light passes through one layer of tissue into another. These properties determine the patterns of light propagation and dispersion, based on which the Monte Carlo results cannot be used to determine the total energy accumulation in different neural tissue regions.

For each specified power density, wavelength, and skin colour the Monte Carlo algorithm was repeated over ten iterations. Previous literature has shown three prominent wavelengths 670 nm, 810 nm, and 1064 nm, which have been modelled in this study. Additionally, common optical power set values range from 100mW/cm^2^ to 300mW/cm^2^, which have also been modelled. To our knowledge, this is the first study to consider the effects of skin colour, and skin pigmentation with PBM. “Caucasian”, “African” and “Asian” skin colours were evaluated with all three wavelengths.

### 2.2 Simulation software

We used Monte Carlo Extreme (MCX), which was developed in 2009 by the Department of Bioengineering at Northeastern University [29]. This project utilized MCX to model the light dissipation of photobiomodulation in the Colin27 head model. The Colin27 atlas was generated as a stereotaxic average of 27 T1-weighted MRI scans of the same individual, linearly registered to create an average with a high signal-to-noise ratio (SNR) developed by the Montreal Neurological Institute (MNI) in 1998 [30]. The Colin27 contains approximately 56.9 million voxels, with a voxel size of 0.5 x 0.5 x 0.5 mm^3^, with each voxel corresponding to a specific tissue type. The MCX simulated normalized energy density of 1 J was modelled with 5e^8^ photons transported through multiple layers of media, including air and tissue. In the simulated scenario, the stimulated spot size is theoretically represented as infinitesimally small. However, in real-world applications, the practical dimension of the disk would range from 0.5-5cm. The 5×10^8^ photons are propelled from the source stochastically, interacting with the various media through absorption, scattering and transmission. As the photon travels, energy is lost according to the Beer-Lambert law, and deposited in the voxel from which it departs, generating a three-dimensional energy deposition map.

### 2.3 Simulation approach

#### tPBM vs. iPBM

Most commonly, PBM is administered through transcranial PBM (tPBM) stimulation by Light-Emitting Diodes (LEDs) or single source laser. This study will position a single source laser between the left and right hemispheres. The default MCX single source optodes sizing, the laser irritation site is the size of one square voxel, or 1 mm^2^. The optode source was positioned for transcranial and intranasal simulation, with the 5×10^8^ incidental photons used in each case, applied over 5 ns. The energy deposition for the laser source is by default normalized to 1 J. The optode location positions are shown in Figure 1a and **1b**.

**Figure 1.**
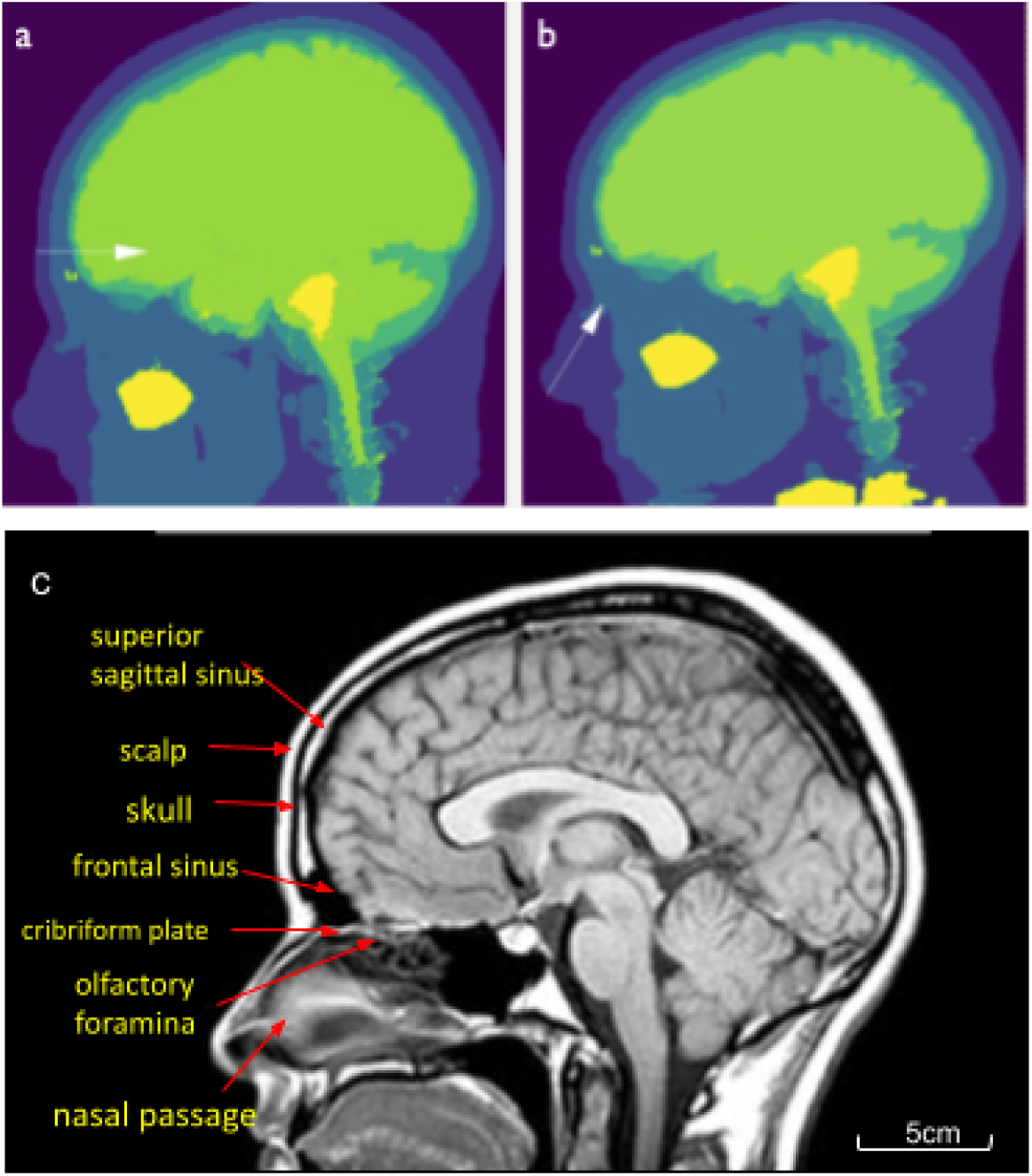
Sagittal slice optode positioning for (a) tPBM transcranial and (b) iPBM settings overlayed onto the Colin27 head atlas. The arrow indicates the position and direction of the flow of photons. Figure 1 (c) highlights the anatomical regions of the brain that both tPBM and iPBM incoming photons must penetrate.

In the tPBM setup, for the light to reach the cortex, it needs to first penetrate the scalp, the skull bone, and the subarachnoid space (containing cerebrospinal fluid (CSF)). In the iPBM set up, instead of the skull, the light penetrates the cribriform plate, which is a part of the ethmoid bone, an unpaired and porous cranial bone that contains many olfactory foramina through which olfactory nerves penetrate the nasal cavity. Whereas the frontal skull is 5-6 mm thick, the cribriform plate is only about 1 mm thick.

iPBM is an emerging stimulation approach that has been shown to improve many brain disorders, including dementia [10]. It is hypothesized that iPBM can project to the hippocampal cortex [31]. Previous literature also suggests that iPBM, due to its access to the brain through the thin and porous cribriform plate, can penetrate deeper into the brain compared to tPBM [26]. iPBM has shown to be a beneficial treatment option for traumatic brain injuries (TBI), sleep disorders and dementia [2], [31].

#### The effect of optical power density

The laser power density has a direct impact on the overall energy accumulation in targeted areas of the brain. The optical power of a PBM light source is measured in watts (W), when accounting for the area of the applied light, this parameter is restructured as optical power density (OPd), which is measured in units of mW/cm^2^ [32]. An average emitted optical power density from an LED or laser of 250-300 mW/cm2 is common in in-vivo human research. The skin-exposure safety limit of near-infrared lasers ranges up to 500 mW/cm^2^ for Classes 3B and 4 lasers, depending on the wavelength and type of laser [33]. As PBM research is still limited, researchers have been experimenting with different ranges of optical power density. A recent 2019 publication reviewed optical power densities ranging much higher than the safety limit, from 200-700mW/cm^2^ in animal studies. This type of research aims to look at heat as a mechanism of action to elicit pain reduction and anti-inflammatory action [33]. This study proved that low levels of optical power were sufficient in obtaining beneficial results without brain heating or damaging effects.

The Monte Carlo Extreme software assumes a normalized incidental total light energy of 1 J. However, to translate the results to more realistic OPd values seen in the PBM literature, including 100, 200 and 300 mW/cm^2^, we calculated power scaling factors for the MCX results. Total incidental optical energy equates to the optical power density multiplied by application time. To scale the normalized energy deposition as quantitative per the actual applied power density to be simulated, the default 1 J incidental energy was divided by a circular area of 1 cm^2^, and the optical power density of the default MCX simulation was calculated using an application time of 60 seconds. Thus, the default power density was 16.7 mW/cm^2^. A scaling factor was then generated and multiplied to the default optical power density and energy values, to scale the MCX results to prominent OPd in PBM literature, namely 5, 7, 9 mW/cm^2^ for iPBM, and 100, 200 and 300mW/cm^2^ for tPBM. The energy deposition (E) over the 60 s window based on each power density level is then normalized by the volume of a given ROI to produce the energy deposition in terms of J/cm^3^.

#### The effect of wavelength

The penetration of light is dictated by wavelength. This parameter determines the distance of which light can travel through the brain. In general, a shorter wavelength, usually in the visible range between 635-700 nm (red light), has shorter penetration than a longer wavelength in the near-infrared range (810-1070 nm) [33]. Literature has suggested that 620, 680, 760 and 820 nm correspond to the absorption spectra of these centres. Many clinical studies utilize an 808nm laser or 810nm LED, reported by some studies to have the deepest penetration and highest cognitive impact in subjects with impaired cognition [16].

The effect of light wavelength is modelled using known scattering, absorption, anisotropy and refractive index coefficients, summarized in Table 2. We obtained these values for 670nm, 810nm and 1064nm, which are commonly used wavelengths in the PBM literature, and which have previously been investigated in simulations [34]. The values are used in Cassano et al [26]. When comparing across wavelengths, a default power density of 100 mW/cm^2^ and 5mW/cm^2^ were assumed for tPBM and iPBM, respectively. A power scaling factor was calculated as described earlier.

**Table 2.** Optical property coefficients. (a) Coefficients for each wavelength, without considering the contribution of skin melanin. (b) Coefficients listed for each skin colour category. CSF = cerebrospinal fluid, µ_s_ = scattering coefficient, µ_a_ = absorption coefficient, n = refractive index coefficient and g = anisotropy coefficient.

| a) |  | Tissue Type |  |  |  |  |
| --- | --- | --- | --- | --- | --- | --- |
| Wavelength | Parameter | Scalp | Skull | CSF | Grey Matter | White Matter |
| | $\mu_s$ | 22.65 | 10.82 | 0.091 | 8.4 | 40.1 |
| **670nm** | μa | 0.05625 | 0.0208 | 0.0004 | 0.02 | 0.07 |
|  | n | 1.37 | 1.37 | 1.37 | 1.37 | 1.37 |
|  | g | 0.89 | 0.89 | 0.89 | 0.9 | 0.85 |
| **810nm** | μs | 17.82 | 17.82 | 0.091 | 7.3 | 38 |
| | μa | 0.0168 | 0.011 | 0.0026 | 0.028 | 0.092 |
|  | n | 1.37 | 1.37 | 1.37 | 1.37 | 1.37 |
|  | g | 0.89 | 0.89 | 0.89 | 0.89 | 0.87 |
| **1064nm** | μs | 14.63 | 14.63 | 0.091 | 5.9 | 30 |
| | μa | 0.019 | 0.019 | 0.0144 | 0.053 | 0.88 |
|  | n | 1.37 | 1.37 | 1.37 | 1.37 | 1.37 |
|  | g | 0.89 | 0.89 | 0.89 | 0.91 | 0.88 |

| b) |  | Skin Type |  |  |
| --- | --- | --- | --- | --- |
| Wavelength | Parameter | Caucasian | Asian | African |
| **670nm** | μs | 21.21 | 21.96 | 14.93 |
| | μa | 0.018 | 0.0212 | 0.0575 |
|  | n | 1.37 | 1.37 | 1.37 |
|  | g | 0.89 | 0.89 | 0.89 |
| **810nm** | μs | 10.45 | 12.73 | 7.95 |
| | μa | 0.0188 | 0.0225 | 0.0285 |
|  | n | 1.37 | 1.37 | 1.37 |
|  | g | 0.89 | 0.89 | 0.89 |
| **1064nm** | $\mu\text{s}$ | 15.26 | 17.05 | 11.08 |
| | $\mu\text{a}$ | 0.014 | 0.0169 | 0.0275 |
|  | n | 1.37 | 1.37 | 1.37 |
|  | g | 0.89 | 0.89 | 0.89 |

**Table 3.** Parameters of Interest

| PARAMETERS OF INTEREST | SYMBOL | UNITS |
| --- | --- | --- |
| WAVELENGTH | $\lambda$ | nm |
| OPTICAL POWER DENSITY | OPd | $\text{mW}/\text{cm}^2$ |
| SKIN COLOUR | - | Eumelanin $\mu\text{g}/\text{mL}$ |
| ENERGY | E | Joules |

#### The effect of skin colour

This was only considered relevant for tPBM. Differences in skin pigmentation are caused by the amount of melanin production. Melanin produced in the innermost layer of the epidermis is a polymeric pigment that produces hair, eye and skin pigmentation. Broad racial differences can be associated with diversity in skin colour, and the concentration of epidermal melanin is double in darker skin types compared to light-pigmented skin types [35]. When light strikes melanin, the pigment molecules within melanin absorb the photons, and skin with higher melanin content absorbs more light. Melanin serves as a natural photoprotective mechanism, absorbing and dissipating photon energy. In regions with intense sunlight and a heightened risk of exposure to harmful ultraviolet (UV) radiation, individuals tend to produce more melanin, resulting in darker skin, as an adaptive protective measure. This elevated melanin content enhances the absorption of UV light, at the skin level, preventing it from penetrating deep into the skin and causing damage. Conversely, individuals with lighter skin exhibit lower melanin concentrations, rendering their skin more vulnerable to light penetration. The absorption of light by melanin plays a pivotal role in the skin’s response to light, with darker skin proving more effective in absorbing light compared to those with lighter skin. Therefore, in terms of photobiomodulation, skin pigmentation is a crucial parameter to consider.

The scalp consists of the skin (i.e. epidermis, dermis, the subcutaneous tissue) and an inner layer (i.e. galeal, subgaleal and periosteal layers). The skin confers the eumelanin that defines skin colour. To accommodate variations in eumelanin concentrations, we adopted three distinct skin colour categories: Caucasian, Asian, and African, the optical properties of which were measured using diffusion optical spectroscopy [34]. The optical property coefficients were extracted from previous Monte Carlo simulation publications [29], [34] where the scalp absorption values were extrapolated using the following equation. The values are summarized in Table 2b.

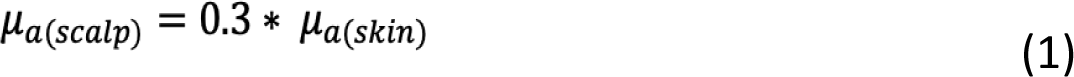

For each parameter setting in each of the categories, the average of the 10 iterations of Monte Carlo simulations was used in all results. We computed the axial energy deposition profile over a 4 cm slab directly adjacent to the position of the light source. We additionally computed percent energy deposition for each condition as a fraction of the incidental laser energy. In-house Matlab scripts [36] were used for compiling multiple iterations of the simulations as well as for visualizing the energy deposition contours and penetration profiles. Finally, based on these profiles, we identified the brain regions of the Colin27 atlas into which the most energy was deposited. These steps were also performed using Matlab.

## 3. Results

### 3.1 Transcranial Photobiomodulation (tPBM)

As highlighted in prior studies, it is important to understand the amount of incidental optical energy delivered to neural tissue. The depth distance between the skin surface and neural tissue is approximately 1-2 cm [9]. Empirically, recent studies applying a 5 W laser, with a 30mm diameter beam size (708 mW/cm^2^) at 808nm on cadaver heads showed that the light can penetrate up to 4 cm from the skin surface. Based on Monte Carlo simulations, Figure 2 displays the modelled energy-deposition profile of the near-infrared light propagation simulated transcranial through the Colin27 brain atlas. According to a sample axial profile derived from the simulations (wavelength = 810 nm), the light energy declines rapidly as it enters the brain (Figure 3). In this case of tPBM, the peak light energy declines sharply over the 5-8 mm that accounts for the skull and sub-cranial tissue layers; light fails to penetrate beyond a depth of 15 mm.

**Figure 2.**
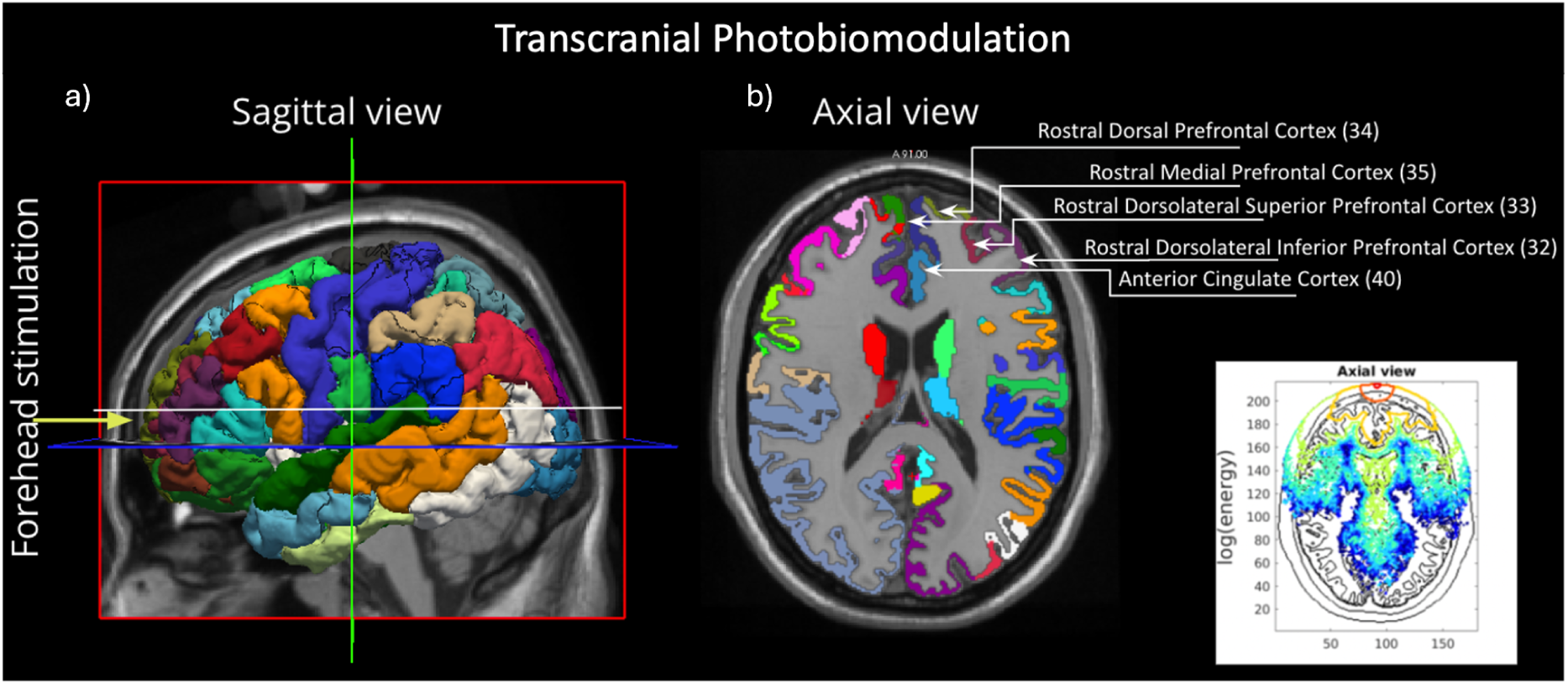
Transcranial PBM irradiation configuration and key regions of interest (ROIs). The yellow arrow represents the laser location, and the white line indicates the axial slice location. The sagittal and axial views of the cortical parcellations are shown in (a) and (b), respectively. The regions receiving the highest energy deposition according to the profile are labelled. The codes correspond to the colour codes used to identify these brain regions in the atlas. Moreover, a sample axial view of the energy deposition profile is shown in the right bottom corner, in which the axes represent the voxel dimensions, and the colour scale represents the log of energy levels.

**Figure 3.**
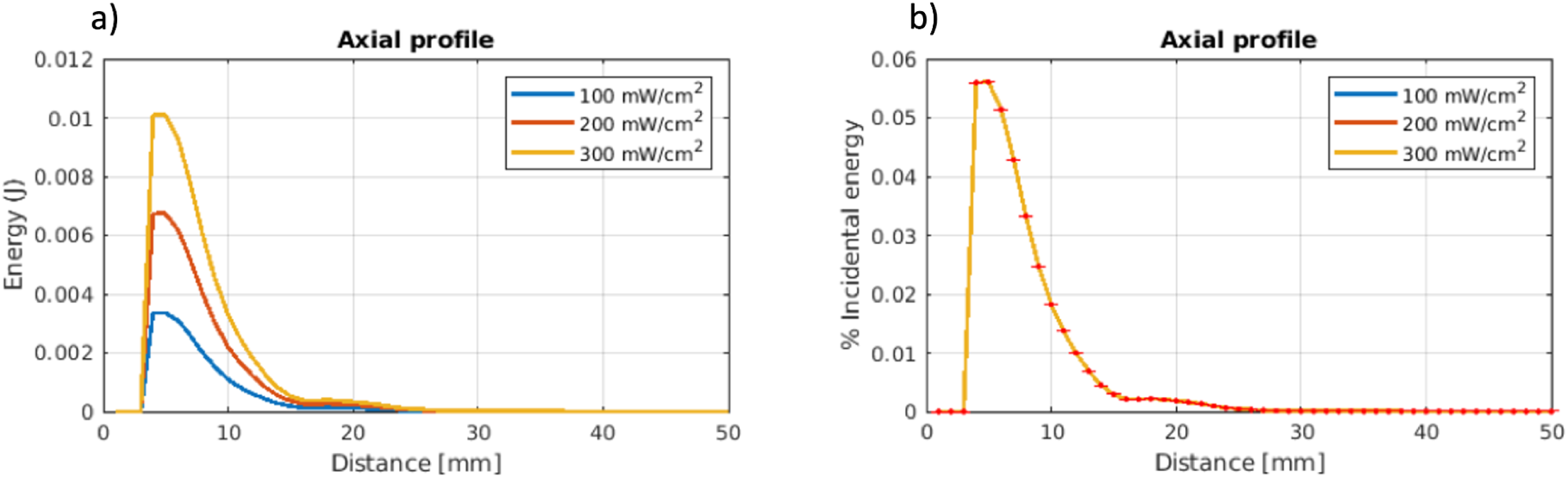
Energy depth penetration profiles for transcranial PBM (tPBM) as a function of power density. A wavelength of 810 nm and the MCX software’s default scalp parameters are used. a) Light penetration depth, based on the rate of energy drop-off with distance in the penetration profile, and subsequent energy accumulation measured posteriorly from the scalp (0mm). b) Percent incidental energy accumulated at the specified penetration depth. Error bars are representative of the standard deviation across the 10 Monte Carlo simulations.

#### 3.1.1 Optical Power Dependence

Our results show a linear response to an increase in energy deposition with OPd, as shown in Figure 4. From this theory, the higher the OPd, the larger energy accumulation in the brain.

**Figure 4.**
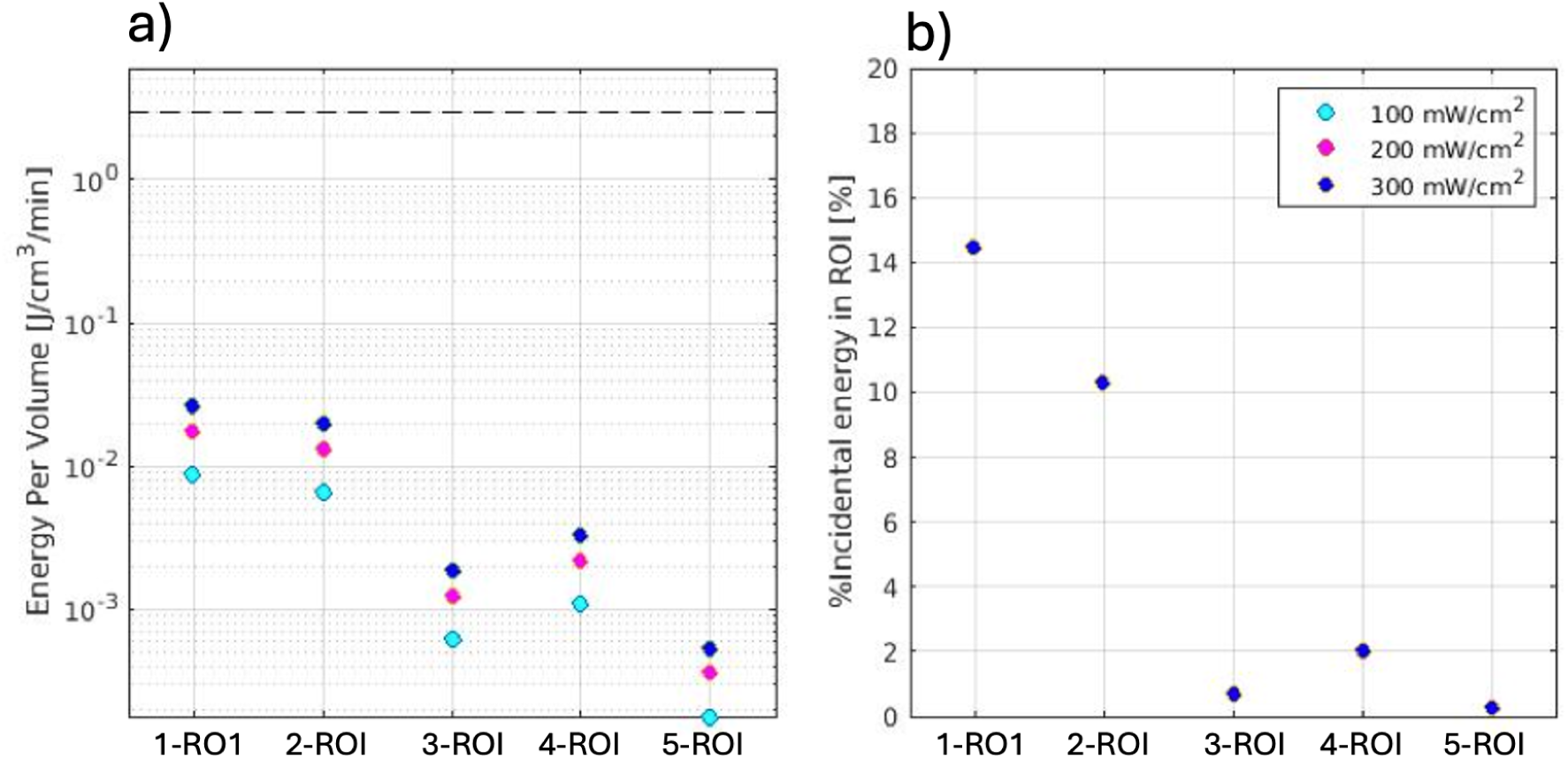
Optical power Plot energy accumulation of transcranial PBM (tPBM) as a function of power density. A wavelength of 810 nm wavelength and the MCX software’s default scalp parameters are used. a) total energy in each region of interest after 1 minute of stimulation (plotted in log scale). b) The percent of incidental energy accumulated in each region of interest, where all three power levels coincide. Note: 1-ROI: Rostral Dorsal Prefrontal, 2-ROI: Rostromedial Prefrontal, 3-ROI: Rostral Dorsolateral Superior Prefrontal, 4-ROI: Anterior Cingulate, 5 -ROI: Rostral Dorsolateral Inferior Prefrontal.

#### 3.1.2 Wavelength Dependence

Our results (Figure 5) show that the 810 nm light had the deepest penetration into the brain for transcranial stimulations.

**Figure 5.**
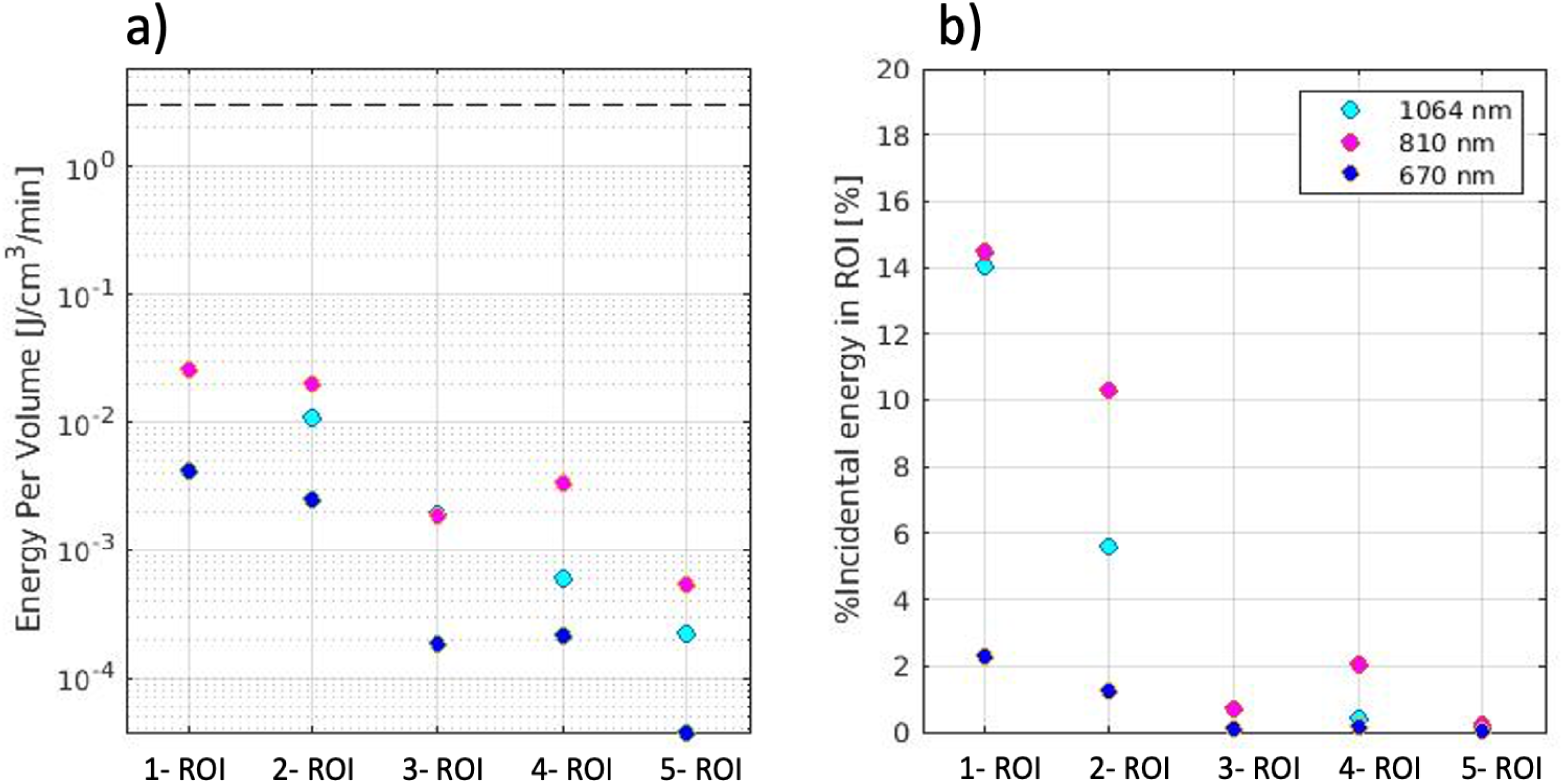
Energy accumulation over key ROIs for transcranial PBM (tPBM) as a function of wavelength. A power density of 100 mW/cm^2^ and MCX software’s default scalp parameters are used. a) total energy in each region of interest after 1 minute of stimulation (plotted in log scale). b) The percentage of incidental energy accumulated in each region of interest, 1064nm percent incidental energy is shown in light blue, which is overlapped by the purple 670nm, in regions of interest 3 and 5. Note: 1-ROI: Rostral Dorsal Prefrontal, 2-ROI: Rostromedial Prefrontal, 3-ROI: Rostral Dorsolateral Superior Prefrontal, 4-ROI: Anterior Cingulate, 5 -ROI: Rostral Dorsolateral Inferior Prefrontal.

#### 3.1.3 Skin Colour Dependence

Taking advantage of the broader skin melanin differences conferred by different races, our results show that those with “Caucasian” skin tone accommodate a higher energy accumulation following tPBM, followed by those of the “Asian” skin tone and lastly, those of the “African” skin tone have a significantly lower energy accumulation. Our results show a wavelength-dependent relationship with skin colour and energy accumulation. The Monte Carlo simulation was repeated for each skin colour and each wavelength, as shown in Figure 6. These results show that the skin-colour dependence also varies with wavelength. The skin-colour dependence in the percentage of energy deposited is highest for 810 nm wavelength, and lowest for 1064 nm. Furthermore, 810nm is associated with the highest percent energy deposition, up to just under 16% for Caucasian skin (Figure 6c)

**Figure 6.**
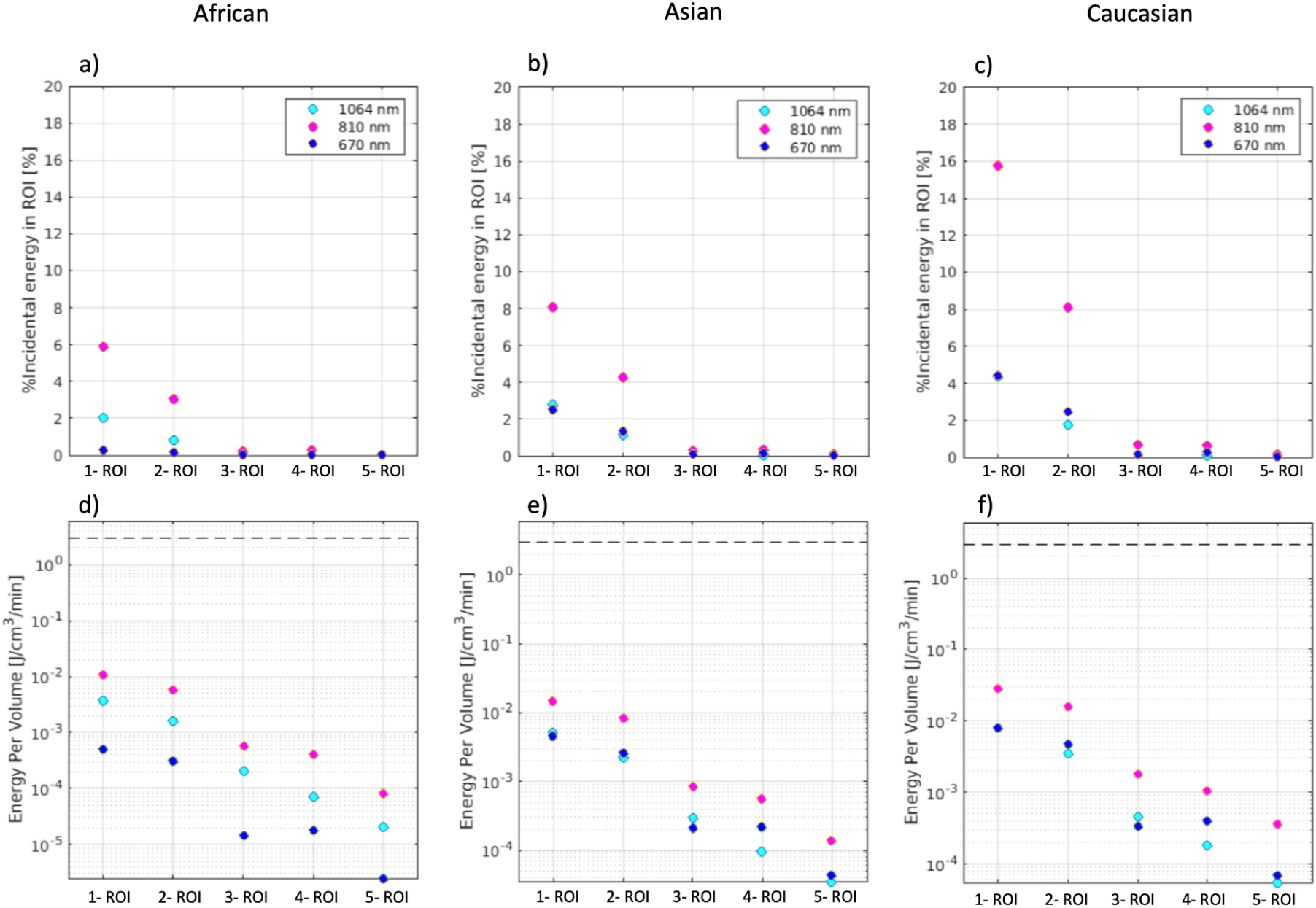
Energy accumulation in key ROIs for tPBM as a function of skin colour type. Transcranial photobiomodulation stimulation, varied skin colour at 670 nm, 810 nm, and 1064 nm wavelengths with an optical power density of 100mW/cm^2^. (a-c) The total energy in each region of interest after 1 minute of stimulation (plotted on log scale). (d-f) The percentage of incidental energy accumulated in each region of interest. Note: 1-ROI: Rostral Dorsal Prefrontal, 2-ROI: Rostromedial Prefrontal, 3-ROI: Rostral Dorsolateral Superior Prefrontal, 4-ROI: Anterior Cingulate, 5 -ROI: Rostral Dorsolateral Inferior Prefrontal.

### 3.2 Intranasal Photobiomodulation (iPBM)

The modelled penetration profile of the near-infrared light propagation simulated intranasally through the Colin27 brain atlas is shown in Figure 7. Our modelling of the propagation of light through the nasal cavity shows that the photon dispersion can lead to a near whole brain response, including the hippocampus.

**Figure 7.**
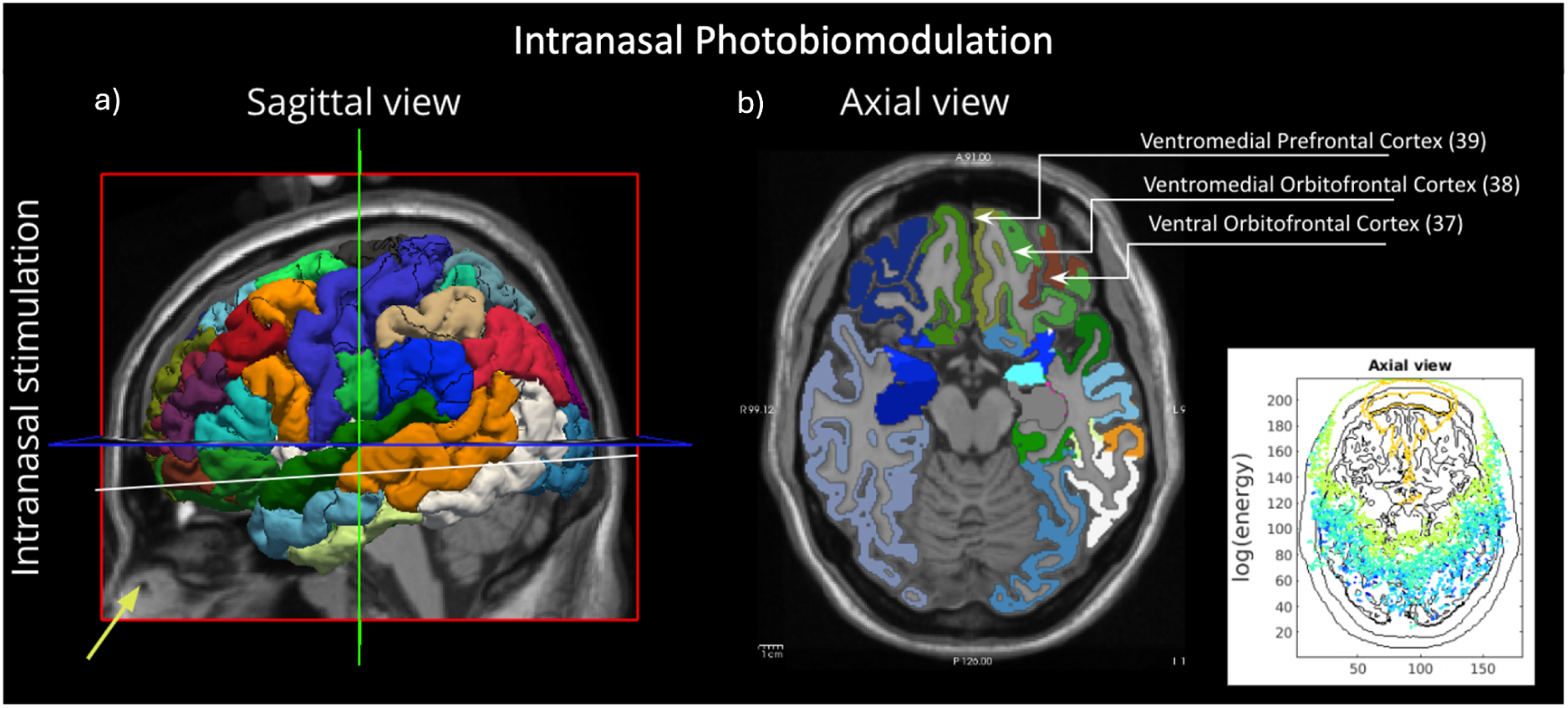
Nasal irradiation configuration and key ROIs. The yellow arrow represents the laser location, and the white line indicates the axial slice location. The sagittal and axial views of the cortical parcellations are shown in (a) and (b), respectively. The regions receiving the highest energy deposition according to the profile are labelled. The codes correspond to the colour codes used to identify these brain regions in the atlas. Moreover, a sample axial view of the energy deposition profile is shown in the right bottom corner, in which the axes represent the voxel dimensions, and the colour scale represents the log of energy levels.

According to a sample axial profile derived from the simulations (wavelength = 1064 nm), the light energy declines rapidly as it enters the nose (Figure 8). In the case of iPBM, the deposited light energy climbs steadily as the light enters the nasal cavity, being deposited in surrounding non-brain tissue. Subsequently, light deposition is reduced at the cribriform plate and in the frontal sinus (about 20 mm from the light source). However, the light deposition subsequently penetrates the thin cribriform plate, reaching about 35 mm from the light source before declining.

**Figure 8.**
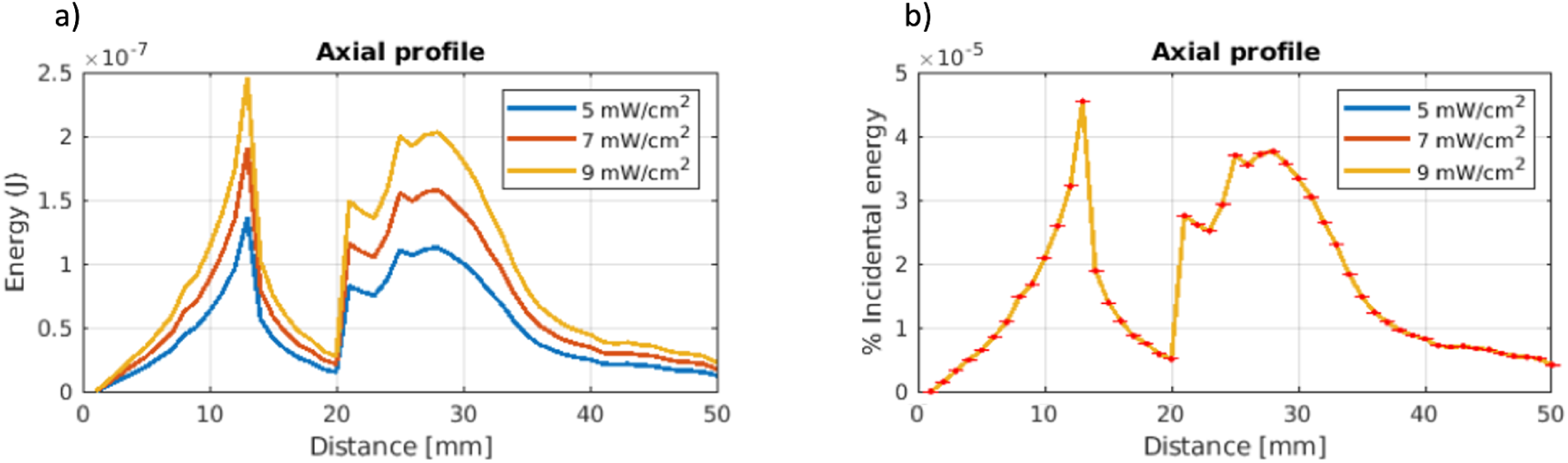
Energy depth penetration profiles for intranasal PBM (iPBM) as a function of power density. Varied optical power densities at 810 nm wavelength. a) light penetration depth, based on the rate of energy drop-off with distance in the penetration profile, and subsequent energy accumulation measured posteriorly from the scalp onwards. b) percent incidental energy accumulated at the specified penetration depth. Error bars are representative of the standard deviation across the 10 Monte Carlo iterations.

#### 3.2.2 Optical Power Density Dependence

Similar to the effect of tPBM on optical power density, our results show a linear response to an increase in OPd, as shown in Figure 9. The 9 mW/cm^2^ power elicited the largest energy accumulation in comparison to the two smaller optical power density values.

**Figure 9.**
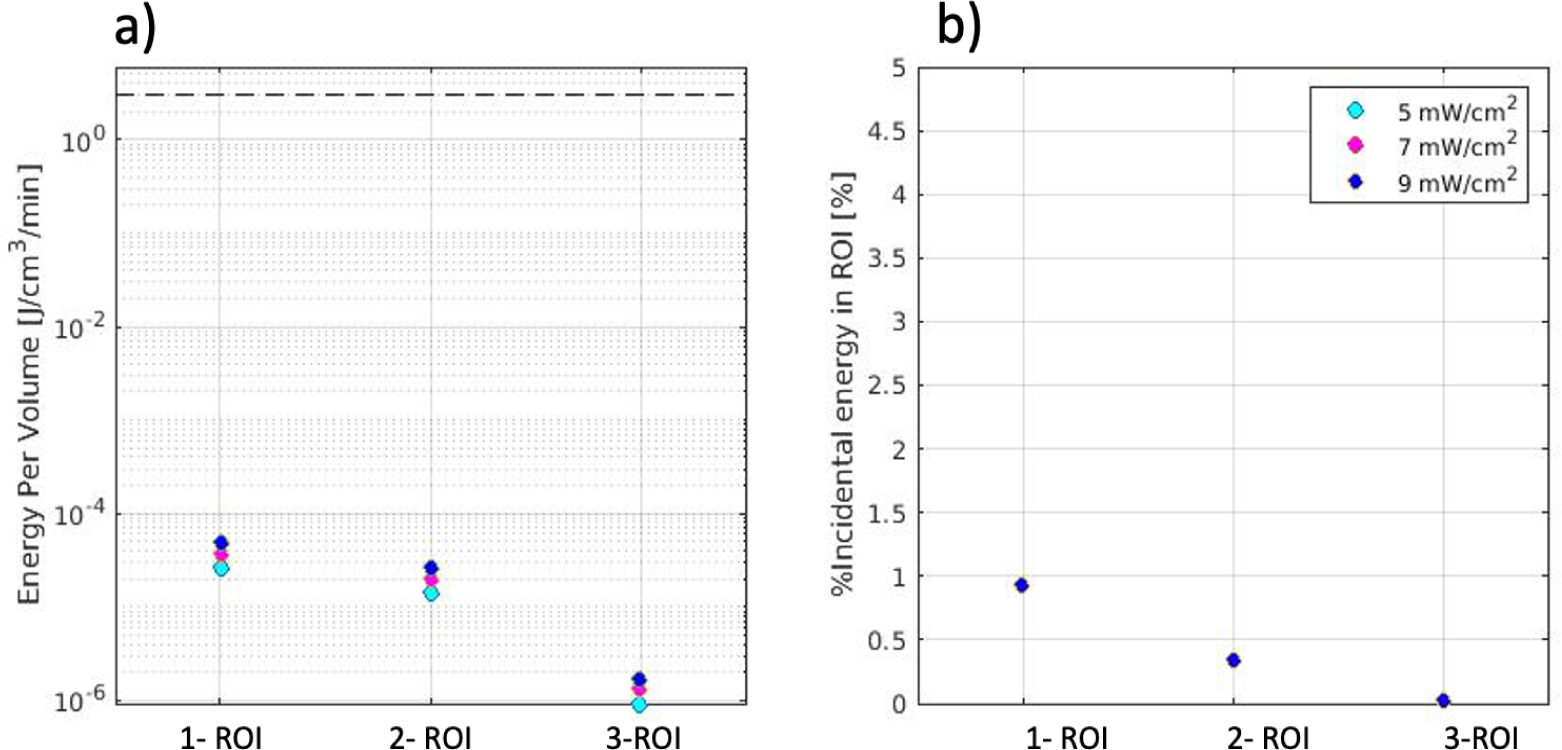
Optical Power Region of Interest Energy Accumulation. Intranasal photobiomodulation stimulation, 810 nm wavelength. a) total energy in each region of interest after 1 minute of stimulation (plotted in log scale). b) percent incidental energy accumulated in each region of interest, where all three power levels coincide. Note: 1 - ROI: Ventromedial Prefrontal, 2 - ROI: Ventromedial Orbitofrontal and 3 - ROI: Ventral Orbitofrontal.

#### 3.2.3 Wavelength Dependence

During iPBM stimulation, the route that the light photons follow does not come into contact with the skull. Unlike tPBM, light transported through the nasal cavity interacts with a significant portion of air when it passes through the frontal sinus. As shown in Figure 8, the energy deposition drops significantly around 15 mm of depth penetration; this drop is hypothesized to be the area of the frontal sinus. Our results (Figure 10) show that the 1064 nm light had the highest percent energy deposition in the brain for both transcranial stimulations.

**Figure 10.**
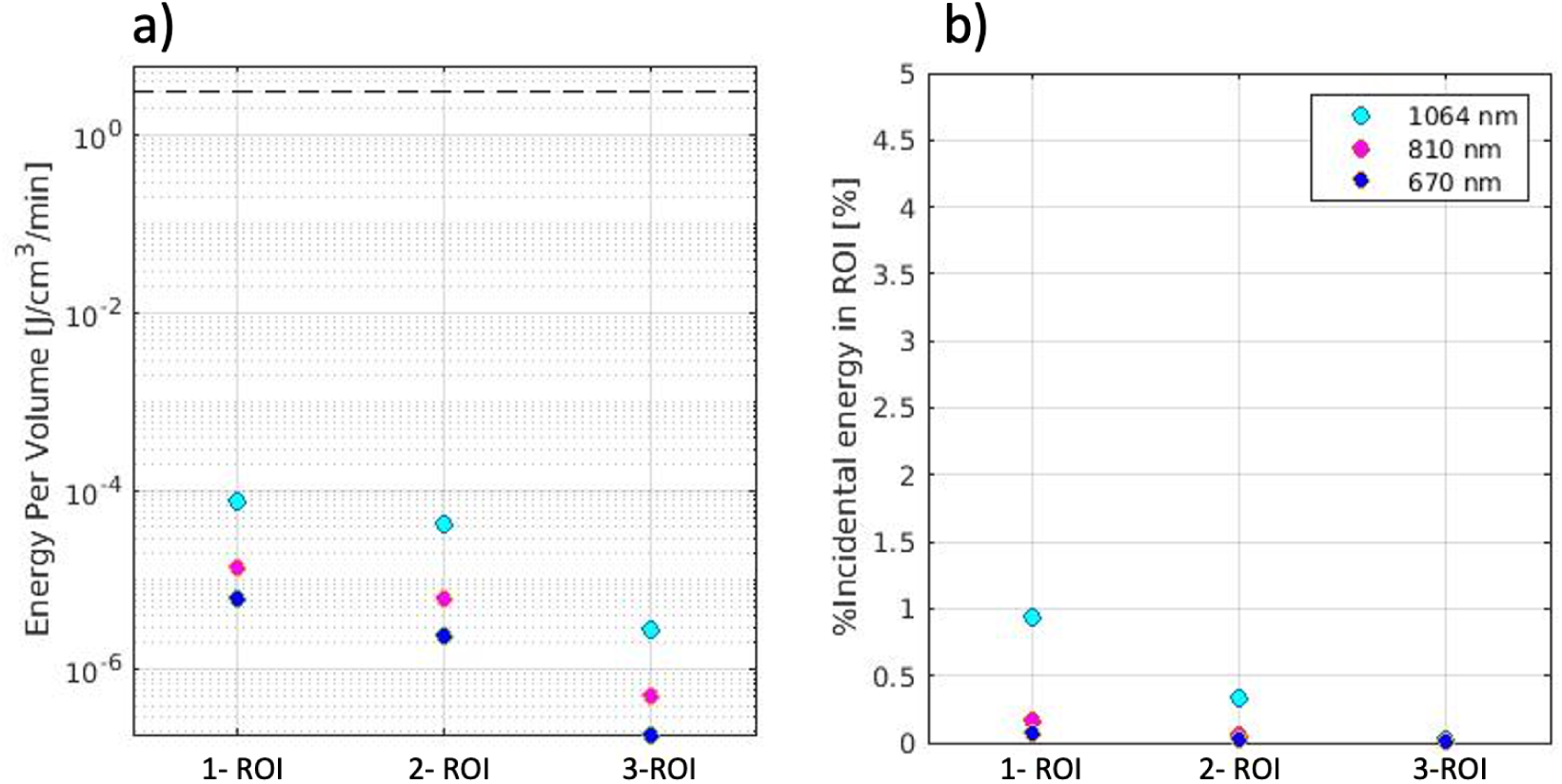
Wavelength dependence of energy deposition in ROIs. Intranasal photobiomodulation stimulation, 5mW/cm^2^ optical power density. a) total energy in each region of interest after 1 minute of stimulation (plotted in log scale). b) The percent of incidental energy accumulated in each region of interest. Note: 1 - ROI: Ventromedial Prefrontal, 2 - ROI: Ventromedial Orbitofrontal and 3 - ROI: Ventral Orbitofrontal.

## 4. Discussion

The Monte Carlo simulation method was first developed to improve decision-making under uncertain conditions [37]. To this day, Monte Carlo simulations have a variety of applications in many mathematical and scientific fields. Assigning a sporadic random value to an uncertain variable in a problem and calculating the result numerous times is the basis behind this mathematical computation. For many linear and complex problems, this simulation technique is considered the gold standard when accounting for unknown variables. However, additional consideration is required when using this technique to model computations related to the human body, especially to that of the brain.

The brain is the most complex organ in the human body [38], consisting of neural tissue, neurons (nerve cells), and glial cells, the brain works as the control system to the human body. Over the last decade, many publications have worked to show the impact of light penetration into the brain through Monte Carlo simulation. To properly model this complex light delivery, several parameters are required, including scattering, absorption anisotropy and refractive index coefficients. These four measurements are provided for each tissue type of the brain: scalp, skull, cerebral spinal fluid, grey matter and white matter. This simulation technique models the linear path of light through each tissue, based on the calculated coefficients. The results of this paper show the effect of multiple variables, including wavelength, optical power density, skin colour, and stimulation location. In this paper, we highlight the distinctions between transcranial and intranasal placements, showing the trajectory of light penetration into neural tissue from forehead and nostril laser placements.

### 4.1 Wavelength

Wavelength determines the distance of which light can travel through the brain. A shorter wavelength, usually in the visible range between 635 - 700 nm (red light), is said to have shorter penetration than a longer wavelength in the near-infrared range (810 - 1070 nm). Monte Carlo simulation is the most common tool for modelling the effect of wavelength on light penetration into the brain. This work shows the effect wavelength has on overall tissue penetration and energy accumulation.

In terms of depth penetration, for tPBM, the 810 nm wavelength produced significantly higher incidental energy deposition than all other wavelengths investigated herein. For iPBM, on the other hand, the 1064 nm wavelength produced the highest energy accumulation. The difference between the modelled penetration profiles for tPBM and iPBM derive from the difference in optode source positioning and the tissue types that they penetrate. With a forehead tPBM the light must first penetrate through the scalp, then skull, followed by CSF, before reaching the neural tissue. The advantage of the 810 nm wavelength in tPBM may be due to it being more distal to the CSF absorption peak. However, for iPBM, the light passes into the nasal cavity and through the porous and thin cribriform plate and the frontal sinus. Not only is the cribriform plate much thinner than the skull (as mentioned earlier), there is also minimal CSF in the path. Thus, iPBM is optimized by the longer 1064 nm wavelength. Moreover, as shown in the light penetration profiles in Figure 3 and Figure 8, the energy dropoff with increasing distance is significantly steeper for tPBM compared to iPBM. However, when averaged over specific ROIs, tPBM deposited higher energy in the rostral dorsal prefrontal (15% - 810nm, 14% - 1064nm) and rostromedial prefrontal (10% - 810nm, 6% - 1064nm) compared to the maximum energy deposition of 1% of the incidental energy for iPBM in the ventromedial prefrontal ROI (Figure 5 and **10**, respectively). However, we are aware of limitations in the accuracy of tissue optical properties measured for wavelengths near 1000 nm. Research at this specific wavelength is less common compared to shorter wavelengths such as 810 nm and 830nm [39].

### 4.2 Optical Power Density

Optical power density is the energy of light, per unit time, delivered to a specific area. The laser power density has a direct impact on the overall energy accumulation in targeted areas of the brain. To optimize energy accumulation, additional Monte Carlo simulations were run to review the impact of optical power and determine the peak value. The energy accumulation results of the Monte Carlo simulation show a linear response to an increase in optical power density. By modelling the light propagation through the human head and scaling to prominent optical power density values, it dictates the total energy accumulated in specific brain regions and its relationship to the OPd parameter. Although this knowledge is crucial for targeting brain areas, it does not consider the metabolic or functional connectivity effects of the incoming light. Previous literature has suggested that the brain’s response to photobiomodulation is biphasic, such that the CCO response is maximized at an intermediate OPd. However, this observation may hinge on physiological processes that cannot be modelled using Monte Carlo simulation.

The current mechanisms of action of PBM are unclear. Recent literature states that incoming red or NIR light interacts with specific cell photoreceptors which can affect cellular pathways [40]. This relationship has been demonstrated in both in vitro cell cultures and human studies, when investigating the ATP and mitochondrial membrane potential upregulation in response to photobiomodulation. Many studies point to the oxidation of CCO, stating that the incoming light photon dissociates an inhibitory (NO molecule allowing the CCO to be oxidized and form ATP). Additionally, the dissociated nitric oxide molecule is a known vasodilator, therefore causing an increase in blood flow due to dilation of blood vessels. This mechanism of action has been discussed in many publications over the past several decades [41], [42]. However, more recently, in-vitro studies have shown that CCO has no impact on ATP production, that cells without CCO present still have an upregulation of ATP following NIR light administration [43]. Other areas of interest include Ca^2^+ channels, K+ channels and reactive oxygen species (ROS). With these uncertainties in PBM mechanisms, the Monte Carlo method is beneficial for modelling the light penetration through the multi-layered tissue of the human head. However, the metabolic and hemodynamic effects remain indistinguishable in this model.

### 4.3 Skin Colour

Melanin, specifically eumelanin, is a substrate in the human skin that produces skin pigmentation, it is the main variable in characterizing skin colour. Human skin can have a vast range of melanin levels, ranging from very low in light complexion; Caucasian skin (type I), to very high in black African skin (type VI) [44]; that is, Caucasian skin is generally known to contain less pigmentation than African and Asian skin types. This study takes the first important step of accounting for the effects of skin colour, and skin pigmentation with PBM, only relevant for tPBM. In this study, we show that although 810 nm deposited the highest percentage of optical power in the cortex for all skin colours, Caucasian skin was associated with a higher overall energy accumulation than other modelled skin colours. However, these skin-colour categories are only names that derive from very limited literature and are not meant to be fully generalizable. That is, in each of the skin-colour categories, there is in reality a wide range of melanin levels. Moreover, not only do skin melanin levels vary across races and ethnicities, they also vary with sun exposure, thus forming an important consideration in personalized PBM.

### 4.4 Implications for human PBM

For the future of photobiomodulation research, it is important to evaluate a multitude of stimulation parameters (wavelength, optical power density, frequency, duration, light source) when working to determine an optimal dosage for PBM therapy. This process is not only crucial for optimizing cognitive outcomes but also allows for comprehension of the wide variations observed in PBM outcomes across diverse studies.

In PBM research, it is apparent that many studies use stimulation parameters that vary significantly. This heterogeneity leads to inconsistent results, making it challenging to replicate findings. Therefore, there is a need to standardize the photobiomodulation parameters to compare across studies, improve the reliability of outcomes, and ultimately advance the understanding of the therapeutic potential of PBM.

Additionally, it is important to understand that the Monte Carlo technique inputs the brain as a static object of several tissue types, therefore disregarding the on-going metabolic or hemodynamic processes. To gain a more comprehensive understanding of how PBM influences the living and dynamic brain, in vivo experiments are warranted. Utilizing imaging techniques like functional magnetic resonance imaging (fMRI) could aid in assessing the accuracy of the Monte Carlo method in reflecting real human subjects.

### 4.5 Limitations

Our study presents several limitations. First, we only probed the forehead tPBM and nostril iPBM positions for the light source. In practice, the positioning of both transcranial and intranasal approaches can vary across studies. An additional limitation of our study revolves around the limited availability of optical coefficient parameters beyond 1000nm. Within the 650 to 850 nm range, melanin absorption dominates, potentially overshadowing weaker absorption traits at higher wavelengths. Additionally, as we delve into the near-infrared (NIR) spectrum past 1000 nm, water starts exhibiting robust absorption characteristics. This added absorption from water presents a competing factor against skin absorption, thereby complicating result interpretation [26], [34].

In addition, our approach did not allow more comprehensive consideration of inter-subject variations. First, we were only able to identify one previous publication that provides documentation on skin attenuation and scattering for various skin types. These values do not define each of the three races (Caucasian, Asian and African), but are rather included as representative values. Skin colour varies widely across each racial category and throughout different seasons, emphasizing the importance of precise measurements rather than relying on broad racial categorizations. Moreover, we did not include sex-related or age-related differences, such as that in cranial thickness [46], into our modeling consideration.

Moreover, there are inherent limitations in the modeling approach, as modeling is almost never truly realistic. For example, the tissue types we modeled were limited by the atlas available in the MCX software. While the main cranial tissue types are included, it was not possible to model tissues such as the mucosae were not included.

Furthermore, our simulation methodology does not enable us to investigate the effects of laser pulsation, blood flow, and other hemodynamic processes. PBM’s impact on dynamic physiological processes is a limitation in PBM Monte Carlo simulations in general. In particular, the flow of neurofluids such as blood and cerebrospinal fluid can alter photon absorption and scattering patterns. The interactions between PBM and cerebral blood flow emphasizes the need for comprehensive simulations that accurately capture the hemodynamic process, thereby enhancing our understanding of the extent to which light can penetrate through the brain.

## 5. Conclusion

Our research represents an important addition to the existing knowledge of light penetration and propagation through the multi-layer tissues of the human head to reach the cerebral cortex. The simulations indicate that even in the cortical region closest to the light, no more than 15% of the incidental energy is deposited into brain tissue. This is higher than previously reported using cadaver heads and skull fragments [16], possibly due to major differences in hydration and other tissue properties between living and cadaver tissues. These physical simulations do not account for dynamic physiological processes that may modulate PBM response in vivo. Thus, these results serve as a starting point to establish the physical baseline of light penetration and factors that modulate it.

## Acknowledgments

We first thank Dr. Qianqian Fang from Northeastern University for his helpful input on using the MCX package. We also thank funding support from the Ontario Centre for Innovation (OCI) and the Natural Sciences and Engineering Research Council of Canada (NSERC). Furthermore, we are grateful for financial donations from Ms. Linda Reed.

